# Exploiting prior knowledge about biological macromolecules in cryo-EM structure determination

**DOI:** 10.1101/2020.03.25.007914

**Authors:** Dari Kimanius, Gustav Zickert, Takanori Nakane, Jonas Adler, Sebastian Lunz, Carola-Bibiane Schönlieb, Ozan Öktem, Sjors H.W. Scheres

## Abstract

Three-dimensional reconstruction of the electron scattering potential of biological macromolecules from electron cryo-microscopy (cryo-EM) projection images is an ill-posed problem. The most popular cryo-EM software solutions to date rely on a regularisation approach that is based on the prior assumption that the scattering potential varies smoothly over three-dimensional space. Although this approach has been hugely successful in recent years, the amount of prior knowledge it exploits compares unfavourably to the knowledge about biological structures that has been accumulated over decades of research in structural biology. Here, we present a regularisation framework for cryo-EM structure determination that exploits prior knowledge about biological structures through a convolutional neural network that is trained on known macromolecular structures. We insert this neural network into the iterative cryo-EM structure determination process through an approach that is inspired by Regularisation by Denoising. We show that the new regularisation approach yields better reconstructions than the current state-of-the-art for simulated data and discuss options to extend this work for application to experimental cryo-EM data.

## 1 Introduction

In cryo-EM single particle analysis, the three-dimensional structure of biological macromolecules is reconstructed from two-dimensional projection images of multiple copies of these molecules in different relative orientations. The requirement to image under low-dose conditions, in order to limit damage to the radiation-sensitive samples, leads to high levels of noise in cryo-EM images and the need to average over many images. Moreover, individual macromolecules, or particles, adopt unknown orientations in the sample and may adopt multiple conformations. The resulting large numbers of unknown variables render the optimimisation problem ill-posed, which combined with the high noise level in the data lead to a challenging reconstruction problem.

The most widely used approach to cryo-EM structure determination to date is based on the maximum-a-posteriori (MAP) estimator. This approach was introduced to the cryo-EM field through its implementation in the RELION program [1, 2]. Other software packages, like Cryosparc [3] and Thunder [4], have since built on the same approach. The MAP approach differs from common alternative approaches in two aspects. Firstly, it marginalises over the unknown orientation (and conformational class) assignments of the particles; and secondly, it uses an explicit penalisation term as a regulariser.

Whereas previously popular software packages typically aimed to find the best orientation and class for each particle, the marginalisation approach rather uses a weighted average of all possible orientations. Marginalisation was introduced to the cryo-EM field by Sigworth [5]. Using simulated data, he showed that marginalisation over the orientations for 2D alignment against a single reference leads to reduced sensitivity to the choice of the starting reference and the ability to align images with very low signal-to-noise ratios. Marginalisation over class assignments was later found to be particularly useful for 2D [6] and 3D [7] classification of experimental cryo-EM images. Since its original conception in Xmipp [6], marginalisation over class assignments has been adopted in many software packages, including RELION [2], Thunder [4], Frealign/Cistem [8], Sphire [9], and Cryosparc [3].

In general, regularisation can be used to prevent overfitting when solving an ill-posed problem. Before the introduction of the MAP approach, cryo-EM software tools would prevent overfitting mostly through heuristic low-pass approaches and Wiener filter approaches [10, 11]. The MAP approach pioneered the optimisation of an explicitly regularised target function by expressing the reconstruction problem in an empirical Bayesian framework. The prior information that underlies the regularisation in this approach is the assumption that rapidly changing density values in real space cryo-EM reconstructions, or high powers at higher spatial frequencies in Fourier space, are unlikely. In other words, one assumes that cryo-EM reconstructions are smooth. By expressing both the (marginal) likelihood function and the prior in Fourier space, a standard *L*_2_-Tikhonov regularisation approach is formulated [12]. Its MAP solution can be obtained through expectation-maximisation [13], where an analytical solution to the maximisation step exists. The resulting algorithm yields a 3D Wiener filter that removes high-resolution noise from the reconstruction, while estimating all necessary parameters from the data [2]. The ability to obtain efficiently filtered reconstructions without the need for user-tunable parameters probably played an important role in the rapid uptake of MAP optimisation in the field [14].

Nevertheless, the smoothness prior seems an information-poor choice when compared to the knowledge about biological macromolecules that has been acquired through decades of structural biology research. For example, we know that the density for the macromolecules in cryo-EM reconstructions is surrounded by flat solvent density and that proteins are made up of amino-acid chains that fold into well-defined secondary structure elements like *α*-helices and *β*-strands. Although a wealth of structural information exists, it has in practice been difficult to hand-craft a formulation of a prior function that can be incorporated into the optimisation algorithm that underlies cryo-EM structure determination.

Machine learning based on deep neural networks, or deep learning in short, can capture this prior information and has recently seen tremendous success in a wide range of computer vision tasks [15, 16, 17]. Convolutional neural networks have been shown to perform equally well or better than conventional state-of-the-art methods in inverse problems for several imaging modalities [18], including de-noising [19, 20, 21], computed tomography [22, 23] and magnetic resonance imaging [24, 25]. Generally, this performance is attributed to the deep neural networks’ ability to efficiently learn statistical correlations about both the noise and the signal through the examples in the training data set.

Regularisation by denoising (RED), is an image-recovery framework that enables the plug-in of various denoisers into a MAP optimisation protocol [26]. The RED framework was originally centered around an explicit expression for a prior that incorporates the output of the denoiser. This framework requires that the denoiser function fulfills two conditions: local homogeneity and Jacobian symmetry. However, most state-of-the-art denoisers, including denoising convolutional neural networks, fail to exhibit these properties, in particular Jacobian symmetry. Despite this, RED has been shown to perform well also with these denoiser functions. Hence, the explicit prior approach cannot explain the behaviour of this framework. In response to these issues, Reehorst and Schniter proposed a new framework, score-matching by denoising (SMD), which showed that RED achieves its performance by approximating the “score” or the gradient of the prior distribution. This approach circumvents the requirement of an explicit prior expression and further does away with the above mentioned conditions on the denoiser function [27].

Here, we propose a cryo-EM structure determination procedure that is inspired by the RED protocol to integrate a convolutional neural network to express prior information. Cryo-EM reconstruction typically uses a vector-valued regularisation parameter to model variations in signal-to-noise ratios across different spatial frequencies. So-called “gold-standard” recommendations in the field are to estimate these through the Fourier shell correlation (FSC) between independently obtained reconstructions from halves of the data set [28]. This estimate typically varies greatly between data sets and throughout different refinement steps. Therefore, we expand RED to account for a vector-valued regularisation parameter that varies throughout the reconstruction and present a theoretical framework, inspired by SMD, to show that this approach is valid in the case of more realistic Gaussian noise. Because we train our convolutional neural network on tens of thousands of cryo-EM reconstructions from simulated data and test it using data that is simulated in the same manner, this work does, not yet, represent a finalised solution to improve cryo-EM reconstruction from experimental data. Rather, it serves as a general proof of principle that learned priors can improve cryo-EM reconstruction. Future research directions, like the exploration of alternative optimisation algorithms and different strategies to design and train the neural networks, are discussed.

## 2 Theory

### 2.1 Mathematical formalization of structure recovery

Structure recovery in single particle cryo-EM is here based on the Bayesian formalization of the problem. The starting point is after the particle picking step where one extracts 2D projection images from the electron micrographs. Each of these shows a single particle. Typically, following a classification step, images of the same macro-molecular structure are selected for further refinement. We may here assume that these 2D projection images are approximately centered (with in-plane 2D translation) w.r.t. the corresponding 3D particle and that minor adjustments remain for an optimal translational alignment.

To formalise the above, let *y*_1_, …, *y*_*m*_ ∈ ℂ^*N*^ denote digitised 2D Fourier transforms of the aforementioned 2D projection images. Likewise, let *s*_1_, …, *s*_*m*_ ∈ ℂ^*M*^ denote the digitized 3D Fourier transforms of the corresponding particles. This 3D Fourier transform acts on the electrostatic potential generated by the particle and its surrounding aqueous buffer. Since all particles represent the same macro-molecule up to an orientation and a translation, we have that *s*_*i*_ *= t*_*i*_ ∘ *r*_*i*_(*x*) where *x* ∈ ℂ^*M*^ is the 3D Fourier transform of the molecule structure, *r*_*i*_ ∈ *SO*(3) is its 3D rotation and *t*_*i*_ ∈ *T* (3) is its 3D isomorphic real-space translation. We define the composite transformation *g*_*i*_ : *t*_*i*_ ∘ *r*_*i*_ and thus *g* ∈ *G* : *SE* (3) belongs to the special euclidean group.

Structure recovery aims to reconstruct *x* ∈ ℂ^*M*^ (the 3D Fourier transform of the structure of the molecule) from 2D data *y*_1_, …, *y*_*m*_ ∈ ℂ^*N*^ when corresponding transormation *g*_1_, …, *g*_*m*_ ∈ *G* are unknown. It is natural to consider data as being generated by a random variable, i.e., there are ℂ^*N*^-valued random variables 𝕪1, …, 𝕪*m* that generate corresponding data. In the Bayesian setting one also adopts the viewpoint that the molecule structure and transformations are generated by random variables. Hence, we introduce a ℂ^*M*^-valued random variable X that generates molecular structures (models) and the *G*-valued random variable 𝕩 that generates particle transformations *g*_1_, …, *g*_*m*_. Data *y*_*i*_ ∈ ℂ^*N*^ generated by the single particle 𝕤*i* := 𝕘*i*(𝕩) is then a single sample of the ℂ^*N*^-valued random variable (𝕪*i* | 𝕩 = *x*, 𝕘*i* = *g*_*i*_) where

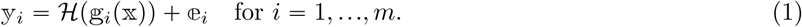

Here, 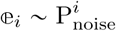 is a ℂ^*N*^-valued random variable representing noise in data and ℋ : ℂ^*M*^ → ℂ^*N*^ is the digitized (linear) model for TEM image formation in frequency space. In particular, *x* ↦ ℋ(*g*(*x*)) = H_*g*_ *x* for some ℂ-matrix H_𝕘_ ∈ ℂ ^*N x M*^ for fixed *g* ∈ *G*. The systematic effects of optical aberrations, like the contrast transfer function (CTF), can be pre-calculated and given during refinement for each image. These effects can be incorporated into, ℋ in which case ℋ would be assigned an *i*-subscript and considered unique for each particle image. However, to simplify the notation, this will be ignored in the following sections.

Assume next that the random variables (𝕪*i* | 𝕩 = *x*, 𝕘_*i*_ = *g*_*i*_) are independent and marginalise over the transformation g ∼ P_*G*_(*g*) using a known (prior) probability density of orientations and translations in *G*. The joint probability density (joint data likelihood) for the entire data set 𝒴 = (*y*_1_, …, *y*_*m*_) ∈ ℂ^*N*^ × … × ℂ^*N*^ conditioned on the model *x* ∈ ℂ^*M*^ is then expressible as

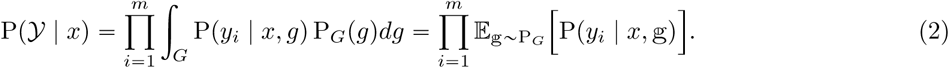

Note here that *x* ↦ P(*y*_*i*_ | *x, g*) for fixed *g* ∈ *G* and *y*_*i*_ ∈ ℂ^*N*^ is given by (1), i.e., it is specified by the matrix H_*g*_ and the noise distribution 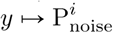.

The Bayesian approach to structure recovery aims to compute a suitable estimator that summarises the (posterior) probability distribution of *x*, given 𝒴. The density of this posterior distribution is by Bayes’ theorem expressible as

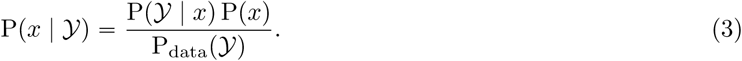

where P_data_ denotes the density for the joint distribution of (𝕪_1_, …, 𝕪_*m*_).

### 2.2 MAP with a Gaussian Prior

The MAP estimator aims to find the model that maximizes the posterior probability (or equivalently its log-posterior probability), i.e., we seek a maximizer to *x* ↦ log P(*x* | 𝒴). From (3) we get that a MAP estimator maximizes *x* ↦ ℒ(*x* | 𝒴) where

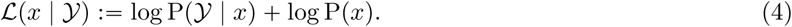

To proceed we need to specify *x* ↦ log P (*x*) (log prior for 3D models) and the joint log-data likelihood *x* ↦ log P(𝒴 | *x*) (joint log-data likelihood).

Assume data in frequency space is corrupted with additive uncorrelated Gaussian noise. More precisely, assume 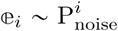 noise in (1) is a complex circularly-symmetric Gaussian with mean zero and diagonal covariance matrix with diagonal vector 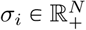 that encodes the frequency-dependent power of noise for each component of the *i*:th image. Then,

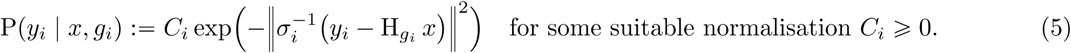

Next, most implementations of MAP-based cryo-EM structure recovery use an uncorrelated Gaussian prior for 3D models with zero mean. Stated more precisely, one assumes

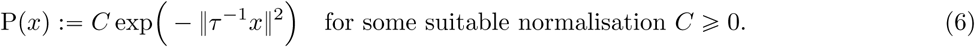

Here, *τ* ∈ ℝ^*M*^ contains the frequency-dependent estimate of the power of the signal in the model. Such a distribution arises from applying the central limit theorem on random distribution of atoms of 3D models, and can thus be intuitively regarded as a minimal assumption criteria about the 3D model [29]. More intuitively, this prior biases the recovery to a smoother real-space representation by suppressing large Fourier components.

A MAP estimator 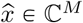 maximizes *x* ↦ ℒ(*x* | 𝒴), so in particular it solves ∇ℒ (*x* | 𝒴) = 0. There is no closed form solution for this equation, so one has to resort to iterative schemes. One example is the Expectation Maximisation (EM) algorithm [13].

For a data likelihood as in (5), the EM method generates iterates {*x*^(*n*)^}_*n*_ ⊂ ℂ^*M*^ by computing *x*^(*n+*1)^ ∈ ℂ^*M*^ from data 𝒴 and the previous iterate *x*^(*n*)^ ∈ ℂ^*M*^. Given (4), (2) and (43) in appendix B it can be shown [30] that this is done by solving the following equation:

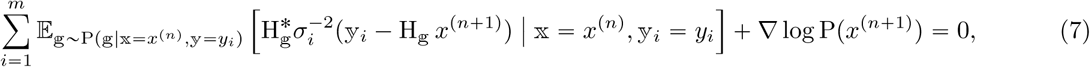

where H^*^ is the adjoint of H with respect to the usual inner product on ℂ^*M*^ (see appendix B) and the expectation is weighted by

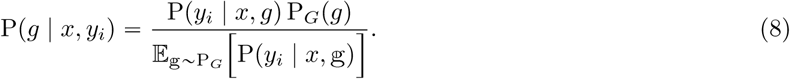

To simplify notation in the following sections we define

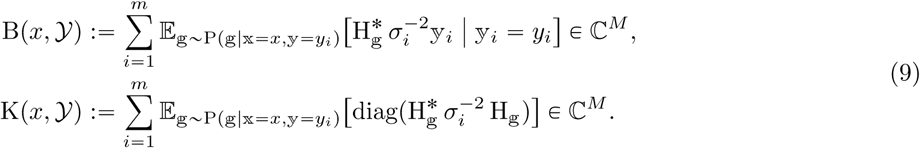

In the above, 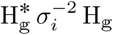 is diagonal due to the Fourier slice theorem and the off-diagonal components are discarded.

Henceforth when multiplying or dividing two vectors of equal length with each other it is to be interpreted as applying these operations component-wise. This allows us to rewrite (7) as

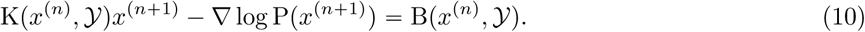

For a prior *x* ↦ P (*x*) that is Gaussian as in (6), we get ∇ log P (*x*) = −*τ*^−2^*x*, so one can solve (7) (perform the M-step) analytically. Which has a closed form solution, generating the following scheme

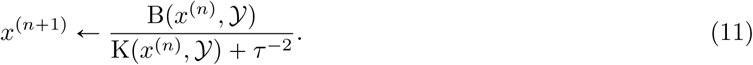

Additionally, we set *σ*_*i*_ ℝ^*N*^ in (9) and the regularisation parameter *τ* ∈ ℝ^*M*^. One common approach in line with the Bayesian viewpoint is to assume these are generated by random variables and then either marginalise over them or use estimates that are updated at every iteration *n*. As an example, the estimate of the regularisation parameter *τ* is commonly based on the radially averaged Fourier shell correlation (FSC) calculated by comparing 3D models reconstructed from two independent halves of the data set [2]. More precisely, let *x*_1_, *x*_2_ ∈ ℂ^*M*^ be two 3D models with associated data sets *𝒴*_1_ and *𝒴* _2_, which are arrays (of equal length *m/*2) with elements in ℂ^*N*^. Next, define

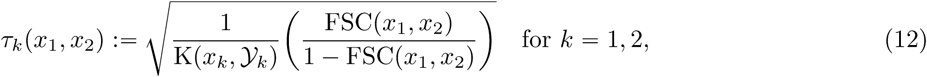

with FSC: ℂ^*M*^ × ℂ^*M*^ → ℝ_+_ denoting the Fourier shell correlation (FSC). To use the above in (11), split the data into two sets *𝒴*_1_ and *𝒴*_2_. Let 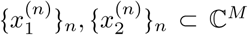 denote the two iterative sequences of 3D models obtained by applying the MAP-EM scheme in (11) on *𝒴*_1_ and *𝒴*_2_, respectively. Instead of using a constant value for the regularisation parameter *τ*, we instead use (12) to adaptively adjust its value based on these sequences, i.e., we replace the fixed *τ* ∈ ℝ^*M*^ in (11) with 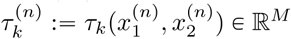 for *k =* 1, 2. Intuitively, this amounts to reducing the regularisation applied in (11) when the FSC increases, which means more signal is permitted to accumulate into each of the two reconstructions.

### 2.3 MAP with non-Gaussian prior

Much of the usefulness of the EM method resides with the ability to perform the M-step in (7) in a computationally feasible manner. This is possible for a Gaussian prior and results in the EM scheme given in (11). Here, we consider priors, *x* ↦ P (*x*), that are “close” to a Gaussian in the sense that the second term below varies slowly with *x*:

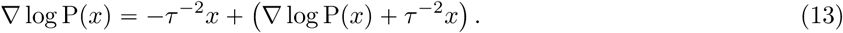

For this class of priors, eq. (10) reads as

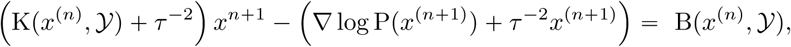

which has the following approximate solution:

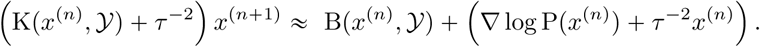

Using this approximation, we can compute an approximate MAP estimator using the scheme:

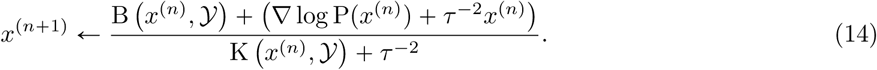

The above requires one to specify the “non-Gaussian” part *x* ↦ ∇ log P (*x*) *τ* ^−2^*x*. One can adaptively set *τ* as in the Gaussian case by splitting data into two parts and using (12). Next, one may consider a data driven approach to learn the “non-Gaussian” part instead of handcrafting it.

The regularization by denoising (RED) method allows us to approximate the gradient of the log-prior using a neural network trained to denoise volumes. The RED prior formula has been shown to accurately estimate the gradient of the prior through score matching by denoising (SMD) [27]. When dealing with independent additive Gaussian noise, RED can be used to integrate an external denoiser into an iterative image restoration protocol [26]. We show in appendix A that the MMSE estimator *f* : ℂ^*M*^⟶ ℂ^*M*^ under Gaussian noise with covariance *τ* ^− 2^ approximates the gradient of the log-prior according to ∇ log P (*x*) *≈ f*(*x*) − *τ*^−2^*x*. However, based on empirical observations, a more conservative approach is to also modulate *f x* with *τ* − ^2^, which would suppress the influence of the denoiser when certainty in the data is high. Hence we propose the gradient log-prior expression

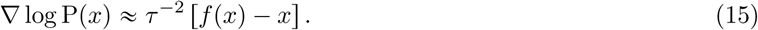

Inserting (15) into (14) gives an approximate M-step for priors using a learned denoiser:

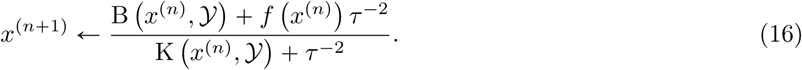

The input to the denoiser in this update formula is *x*^(*n*)^, which lacks the information extracted in the latest E-step. However, both K and B can be evaluated at this stage, hence through them it is computationally feasible to provide the denoiser with the most “up-to-date” information. Additionally, based on empirical observations we introduce a heuristic paramater, *λ*, to tune the influence of the denoiser in the M-step. Conclusively, we propose the following update scheme:

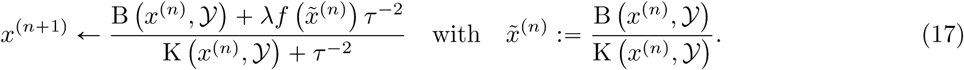

Note that (17) is equivalent to (11) when *λ* = 0, so 0 ⩽ *λ* ⩽ 1 is an empirical parameter that balances the influence of denoiser vs. conventional Gaussian prior. Furthermore,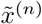 in (17) is equivalent to *x*^(*n*+1)^ in (11) if *τ* ^−2^ = 0. Hence the denoiser acts on the unregularised map that contains information from the most recent alignment of the experimental data, which alleviates the assumptions made for (14).

Further adaptation can be obtained by making use of the fact that RELION is used for the refinement. To do this, we consider an mimimum mean-square estimator (MMSE) 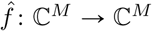 that is trained to “denoise” RELION refinement volumes. More precisely, we consider the following supervised statistical learning problem:

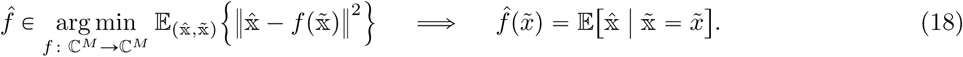

In the above, 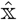 is the random variable that generates appropriately coarsened (low-pass filtered) 3D structures of macro-molecular assemblies from the PDB and 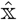 is the random variable that generates RELION refinement structures.

Since we lack knowledge about the joint probability distribution of 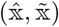, the conditional expectation in the left-hand-side of (18) is replaced by its empirical counterpart given by supervised training data 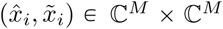 for *i =* 1, …, *n* that are random draws from 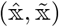, i.e., 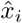 is the true 3D structure derived from PDB (after appropriate coarsening) and 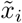 is a corresponding RELION refinement structure. Furthermore, a deep neural network is used to parameterise ℂ^*M*^-valued mappings on ℂ^*M*^, thereby replacing the infinite dimensional minimisation over such functions with a finite dimensional minimisation over the deep neural network parameters. The resulting counterpart of (18) can then be written as the following empirical risk minimisation problem:

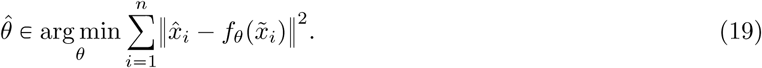

If the network has sufficient model capacity and there is enough supervised training data, then the above yields an approximation to the conditional mean estimator, i.e.,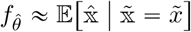.

## 3 Experimental Design

### 3.1 Convolutional Neural Network

Synthetic training data was generated from 543 atomic structures that were downloaded from the Protein Data Bank (PDB) [cite]. All downloaded structures were solved using X-ray crystallography to at least 4 Å resolution; had molecular weights between 40 and 100 kDa; and consisted of a single protein chain. 110 structures also contained nucleic acids. Atomic models were converted to volumetric density maps in the MRC format [cite] using the pdb2map-tool in EMAN2 [31]. In order to obtain simulated projection data sets at different signal-to-noise ratios (SNRs), the intensity values of these maps were multiplied with three different constants: 0.014, 0.012 and 0.010. Next, the density maps were projected onto images of (96 × 96) pixels, with a pixel-size of 1.5Å, and independent Gaussian noise of unity variance was added. The resulting projection data sets had SNRs in the range 0.0003 to 0.0100 and on average 0.0020, which was calculated as the average per-pixel variance of the projection images divided by the variance of the added noise. No attempts were made to model the typical CTF of an electron microscope. Individual projection data sets of 10,000 projections each were generated for the different structures and SNRs using random projection directions that uniformly sampled the orientation space. Each of these projection data sets was then used for standard, unmasked “3D auto-refinement” in Relion [2], which was started from a 30Å low-pass filtered version of the ground truth map. New external-reconstruct functionality in Relion-3.1 (also see below) was used to obtain unregularised half-maps at each of the intermediate iterations of all refinements. In total, this process yielded over 25,000 unmasked 3D density maps at different stages of refinement, which were further augmented with size-preserving rotations.

To verify the validity of the underlying assumptions about Gaussian distribution of noise for the derivations in the theory section we also trained a network using pairs of noise free ground truth and pure Gaussian noise distorted maps. For this purpose we used the estimated frequency space distribution of noise calculated from the Relion intermediate reconstructions. During training, random Gaussian noise was generated in frequency space matching this profile. The noise was then convoluted with a real space mask that dampened the noise levels in the solvent region. However, unless specified the presented results are for the a denoiser trained on Relion intermediate reconstructions. All networks where trained and evaluated on real-space maps. In practice, each map update involves an inverse Fourier transform of 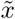 and a subsequent Fourier transform of the denoiser, *f*, before (17) is evaluated.

For this work we chose a U-Net [17] for the denoiser and trained the network using Tensorflow 1.15 [32] in Python. The network was trained with residual learning [19] via ADAM [33] with a mini-batch size of 10 maps for 27 epochs. The U-net consisted of five down- and up-sampling stages that were implemented through max-pooling and transposed convolutional layers [34], respectively. Each stage was made up of two consecutive blocks repeating BN+ReLU+Conv. The input-output pairs for training consisted of the unregularised and unmasked reconstructions from intermediate Relion iterations and their respective noise-free ground truth maps. Here, the coarsening (low-pass filtering) applied to the ground truth (see (19)) was achieved by low-pass filtering it to match the resolution of the corresponding noisy input map. This is done by multiplying the Fourier transform of the ground truth with the FSC of each noisy input map. The FSC is estimated for each map by Relion during the reconstruction of the training data set from the half-maps.

### 3.2 Assessment of the RED approach

Four PDB structures were selected to test the performance of the RED approach (Table 1). The test structures were excluded from the training data set and had a minimum of 8Å RMSD difference for at least 1,000 aligned atom pairs with any other structure in the training data set. The four test structures represent different levels of difficulty for the reconstruction, mostly arising from differences in their molecular weight (MW) and their overall shapes (as projection images of near-spherical structures are harder to align than those with more anisotropic shapes). In order of increasing difficulty the test structures are listed in Table 1, from the elongated, larger structure 4AIL to the smaller, near-spherically shaped structure 4M82. Projection data sets for testing were made in the same way as the projection data sets that were used to generate the training set for the convolutional neural network. Test data sets were generated at four different SNRs by multiplying the original maps from the pdb2map tool by 0.014, 0.012, 0.01 and 0.008, which yields the average SNRs (*i*)0.0038, (*ii*) 0.0028, (*iii*)0.0019 and (*iv*) 0.0012, respectively.

**Table 1:** Characteristics of structures in the test data set. Compactness is expressed as the ratio between the smallest and the largest diameter of the molecule. Relative image SNR is expressed as the per-pixel average variance of signal over noise relative to that of the structure with the maximum SNR (4AIL).

|  | 4AIL | 4BB9 | 4BTF | 4M82 |
| --- | --- | --- | --- | --- |
| Molecular weight ($kDa$) | 96 | 70 | 54 | 46 |
| Relative image SNR | 1.00 | 0.52 | 0.27 | 0.26 |
| Compactness | 0.42 | 0.64 | 0.62 | 0.74 |
| α-helical content (%) | 38 | 49 | 46 | 39 |
| β-strand content (%) | 21 | 10 | 10 | 12 |
| Loop content (%) | 36 | 41 | 44 | 49 |
| Nucleotide content (%) | 5 | 0 | 0 | 0 |

The single-pass performance of the denoiser can be examined by applying it once to unregularized maps and evaluating the ratio between the *L*^*p*^-difference to ground truth from the denoised map and the unregularized map. We evaluated the average of this ratio (using both *L*^1^ and *L*^2^) for the entire training data set, as a function of the estimated nominal resolution of each map.

Standard, unmasked Relion 3D auto-refinements, with updates based on (11) were compared to refinements with updates based on (17). Again, all refinements were started from initial models that were obtained by 30 Å low-pass filtering of the ground truth map and used the same refinement parameters. The results of both types of refinements were compared based on the reported half-map FSC and the FSC against the ground truth and the angular error relative to the true projection angles.

### 3.3 Implementation details

Instead of implementing the update formula in (17) directly in the C++ code of Relion, we created an external reconstruction functionality in its refinement program. When using the `--external_reconstruct` commandline option, the `relion_refine` program will write out MRC files containing B (*x, y*)and K (*x, y*) for both half-maps at each iteration step, together with the current estimate for FSC and *τ* ^2^. The program will then wait until reconstructions with a specified filename for both half-maps exist on the file system, read those maps back in, and proceed with the next iteration of refinement. This functionality was coupled to a script in Python to perform the update formula in (17) using the pre-trained denoiser model. The Python script for external reconstruction using the proposed RED protocol, together with the architecture and the weights of the convolutional neural network that was used for the presented results may be downloaded from github (https://github.com/3dem/externprior).

## 4 Results

### 4.1 Performance of the denoiser

We first tested the effects of applying the denoiser on individual, intermediate, unregularised reconstructions from a standard Relion refinement (Figure 1 A & B). After the first iteration of the refinement of the 4AIL projection data at SNR (*i*), the unregularised half-reconstruction contains only lower resolution components and exhibits notable rotational smearing. The latter is often observed in low-resolution refinement intermediates and is caused by uncertainty in the angular assignments. Application of the denoiser reduces the rotational smearing and brings the map closer to the true signal, as confirmed by visual comparison and the FSC with the ground truth map.

**Figure 1:**
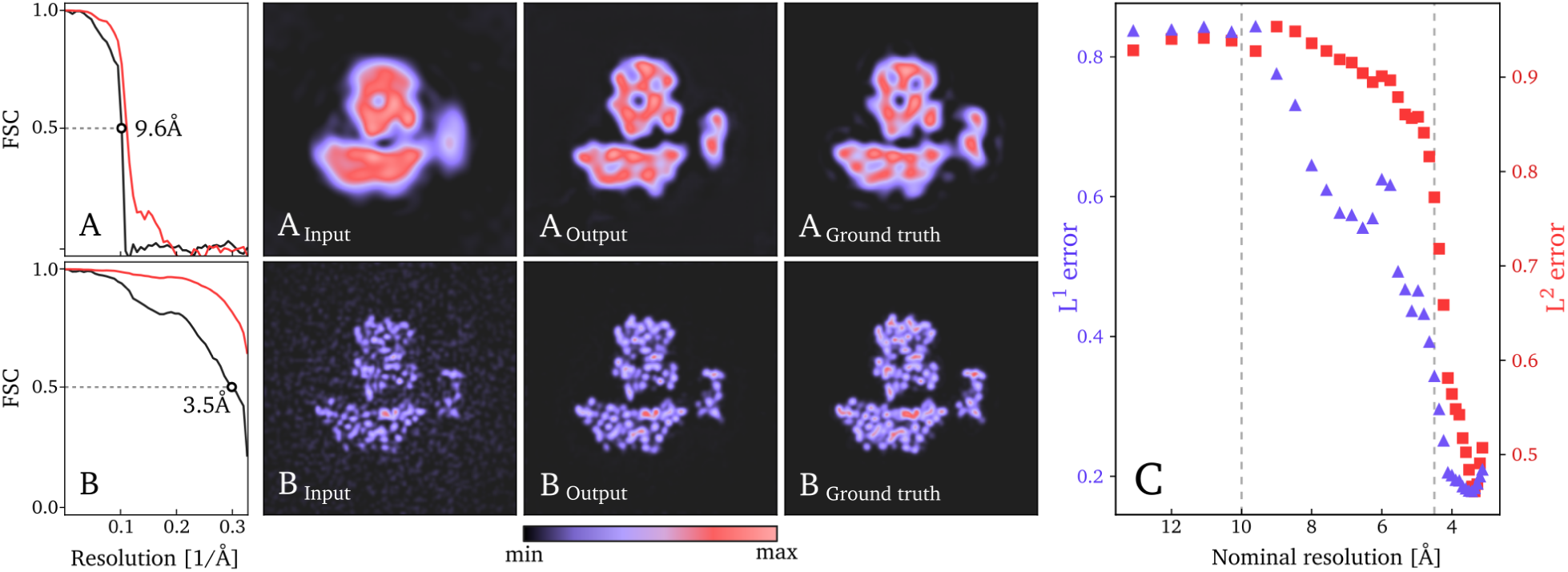
Single-pass denoising performance results, at low (**A**) and high (**B**) resolution, showing central slices of unregularised input map, output from the denoiser and ground truth for reconstructions of a structure from the test data set (PDB-ID: 4AIL). Color-scale spans between minimum and maximum density values in each slice. FSC curves are showing correlation with ground truth before (black) and after (red) denoising. The resolution at *FSC =* 0.5 is shown for each curve. **C** shows the average ratio between the *L*^*p*^-difference from ground truth to the denoised map and the unregularized map, for the entire training data set, as a function of the nominal resolution.

At iteration 16 of the same refinement, the unregularised map contains higher resolution components, but noise components are clearly visible in the volume surrounding the protein, which should be devoid of features. The denoiser efficiently removes the latter and, again, both visual comparison as well as FSC calculation with the ground truth map confirm that the denoised map is closer to the ground truth signal than the unregularised map.

The average ratio between the difference to ground truth from the denoised map and the unregularized map was calculated for *L*^1^ and *L*^2^, at each nominal resolution (Figure 1 C). When the resolution of the input map is worse than 10Å the denoiser fails to produce a significant improvement on average. As the resolution improves beyond 10Å the performance gradually improves and eventually plateaus beyond 4.5Å.

### 4.2 Performance of the RED approach

Next, we tested the performance of the RED approach on projection data sets of the four test structures at four different SNRs. First we run the RED with a denoiser trained on Gaussian noise and compare the results with that for a denoiser trained on Relion intermediate reconstructions (Figures 2). The RED based approach outperforms the standard approach in all refinements performed, as measured by FSCs between the resulting reconstructions and the ground truth maps (solid green and purple lines in Figure 2). This suggests that Gaussian noise can partially explain the distortions. However, the denoiser trained on Relion intermediate reconstructions performs better, confirming that Gaussian noise is an incomplete model of the distortions observed in Relion intermediate reconstructions.

**Figure 2:**
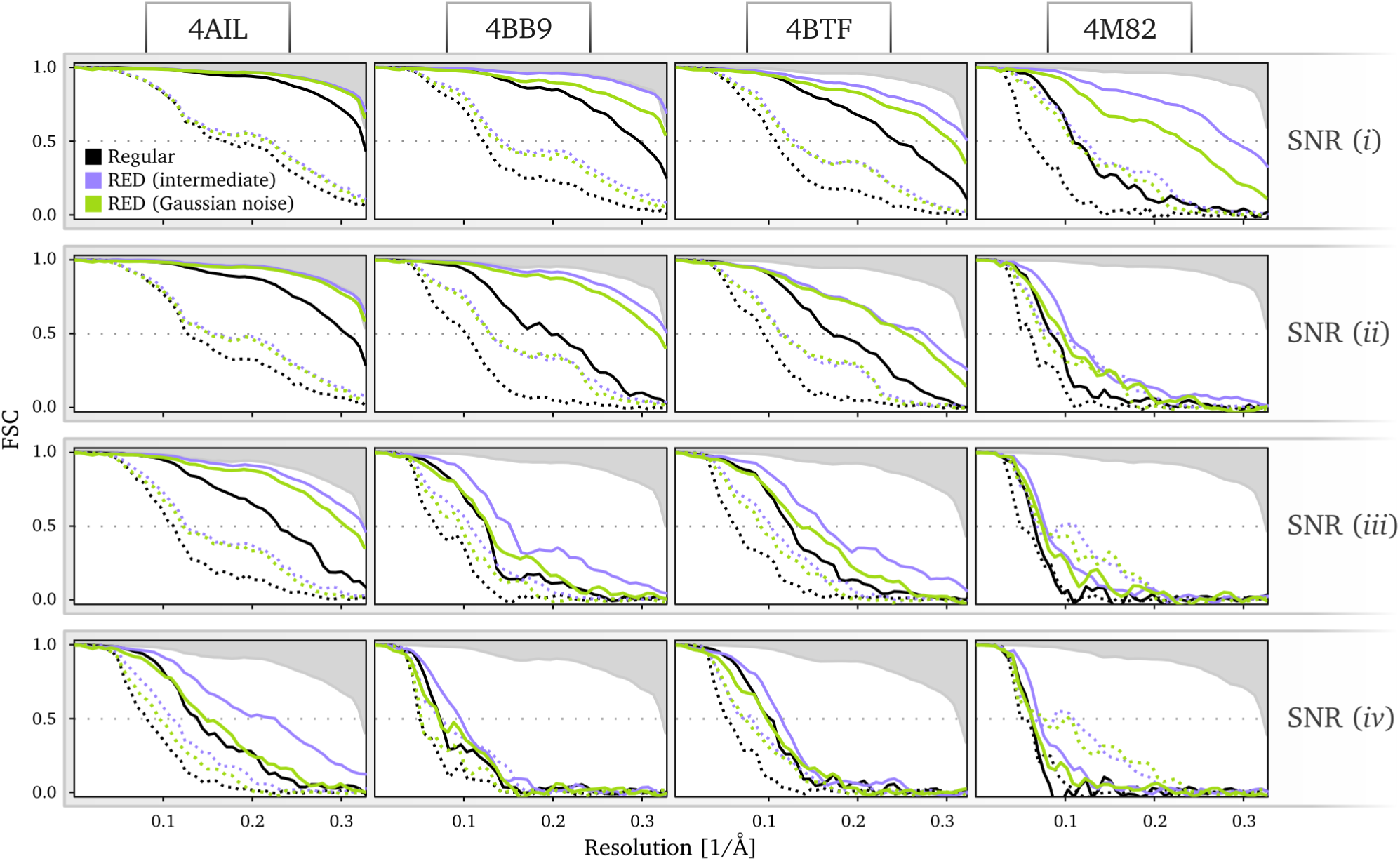
FSCs of reconstructions at four different SNRs (rows) of four structures (columns). Comparing regular Relion (black) with RED, using a denoiser trained on Gaussian noise (green) and on Relion intermediate reconstructions (purple). Dotted lines show the half-map FSC and solid lines show FSC between the regularised map and ground truth. All maps are first multiplied with a solvent mask with a smooth edge before comparison to ground truth. Upper shaded area shows the FSC when model is reconstructed with zero angular error.

Both the RED approach and standard Relion auto-refinement produced a reconstruction with a resolution close to the Nyquist frequency for the easiest test structure, 4AIL, at the highest SNR. Conversely, both approaches yielded a low-resolution solution that was devoid of recognisable protein features for the most difficult test structure, 4M82, at the lowest SNR. Therefore, the range of structures and SNRs of our test data set represents a wide range of scenarios, from easy to hard refinements. The gains in performance tend to be higher at higher SNRs; they are small for the most difficult refinements.

Also measures of reconstruction quality that are reported by Relion without having access to the ground truth, i.e. the half-map FSCs, improve when using the RED approach (dotted green and purple lines in Figure 2). However, for the refinement of 4M82 at the three lowest SNRs, we noticed a severe overestimation of resolution based on the half-map FSCs using the RED approach. A visual examination of the maps (figure 4 B) reveals high-contrast features that are similar in appearance to those of biological macromolecules, but do not correspond to ground truth. This points to the denoiser introducing false high-resolution features into the reconstructions that are shared among the two half-maps. This then leads to an overestimation of the resolution, which is based on the correlation between the half-maps. Since the half-map FSC is the only reliable estimate of resolution when ground truth is missing this issue poses a major problem for the validation of the reconstruction results.

### 4.3 Confidence Weighting

The *λ*-parameter in (17) can be used to fall-back on the Gaussian prior in a scenario where confidence in the performance of the denoiser is lacking. Inspired by the single-pass performance of the denoiser (Figure 1 C) we tested a simple tuning scheme for *λ* to address the issue of resolution overestimation. We will refer to this modification as *confidence weighted RED* (CW-RED).

At nominal resolutions worse than 10Å, as estimated by Relion’s half-map FSC, we assign a full fall-back onto the Gaussian prior, by setting *λ =* 0. This is where the single-pass performance indicates that the denoiser results in little improvement in the reconstruction (see Figure 1C). To avoid impairing the performance at high-resolutions, however, we set *λ =* 1 at nominal resolutions better than 4.5Å, which is approximately where the single-pass performance of the network begins to peak. For any nominal resolution between 4.5Å and 10Å we use a linear interpolation between zero and one.

Using this approach we see that the overestimation of the half-map FSC is significantly alleviated (dotted red lines in Figure 3) without affecting the overall quality of the reconstructions as measured by the FSC with ground truth (solid red lines in figure 3); the errors in the angular assignments (Figure 5); and the visual appearance of the resulting structures (Figure 6 and 7). A visual inspection of the 4M82 map resulting from the refinement with the lowest SNR also suggests that the reconstruction no longer accumulates false structural features with poor correlation to ground truth (Figure 4).

**Figure 3:**
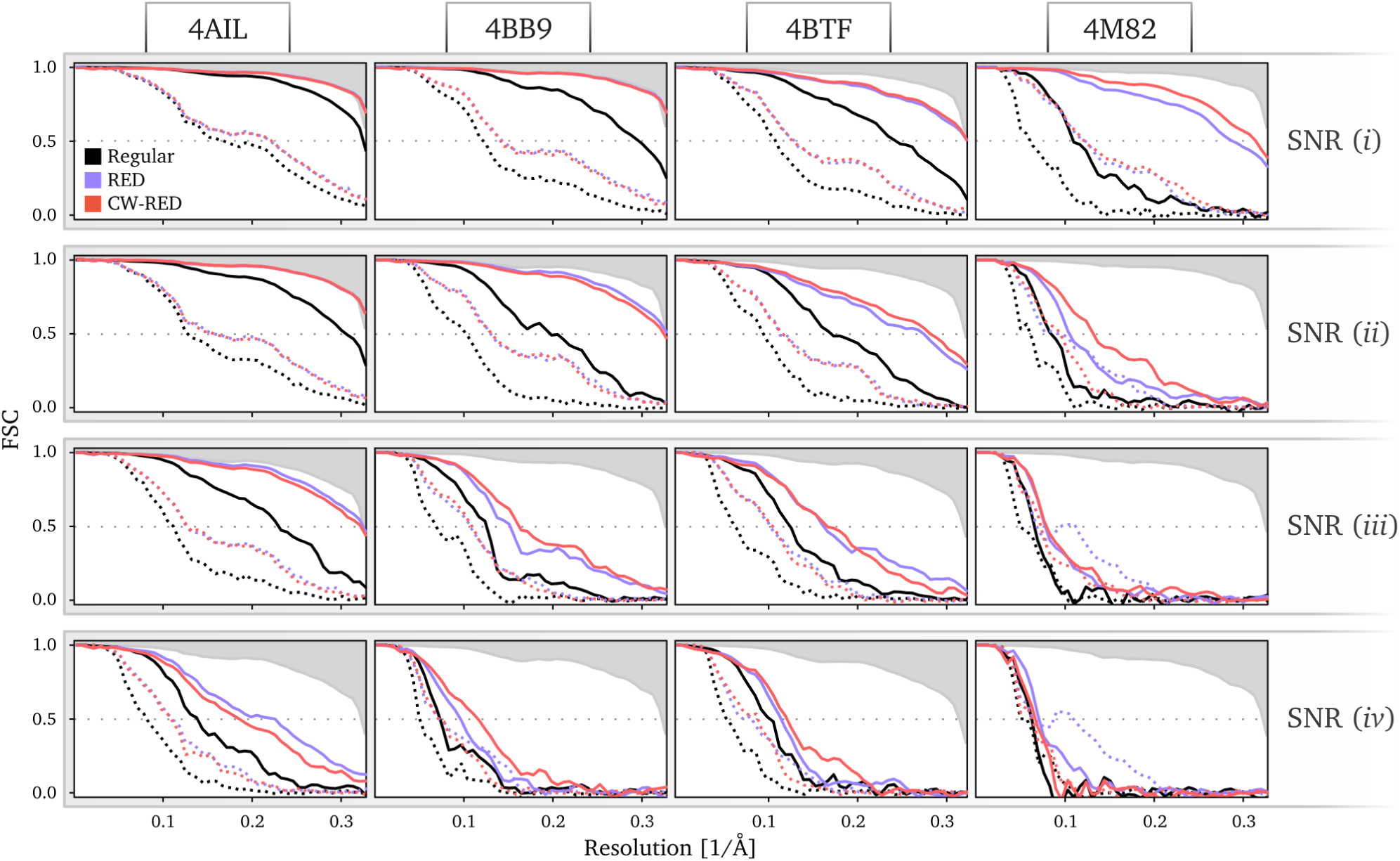
FSCs of reconstructions at four different SNRs (rows) of four structures (columns). Comparing regular Relion (black) with RED (purple) and confidence weighted RED (red), both using denoiser trained on Relion intermediate reconstructions. Dotted lines show the half-map FSC and solid lines show FSC between the regularised map and ground truth. All maps are first multiplied with a solvent mask with a smooth edge before comparison to ground truth. Upper shaded area shows the FSC when model is reconstructed with zero angular error. Note that purple lines are showing the same results as the purple lines in figure 2.

**Figure 4:**
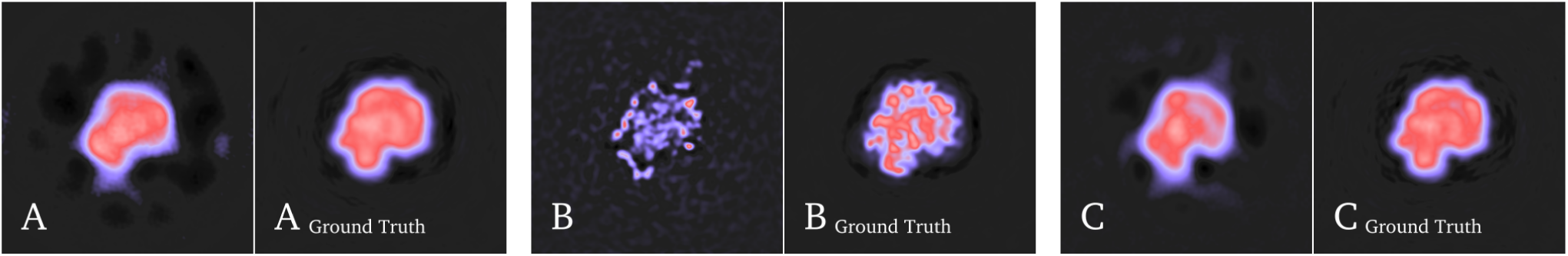
Central slices of reconstruction results of 4M82 at SNR (*iv*) and their respective ground truth. Regular Relion (**A**), RED (**B**), confidence weighted RED (**C**). Each ground truth map has been low-pass filtered to match the estimated resolution of the reconstructed maps by multiplying their Fourier transform with the half-map FSC. The result for RED is showing high-contrast features that do not correlate well with ground truth. This issue is significantly alleviated by confidence weighting.

**Figure 5:**
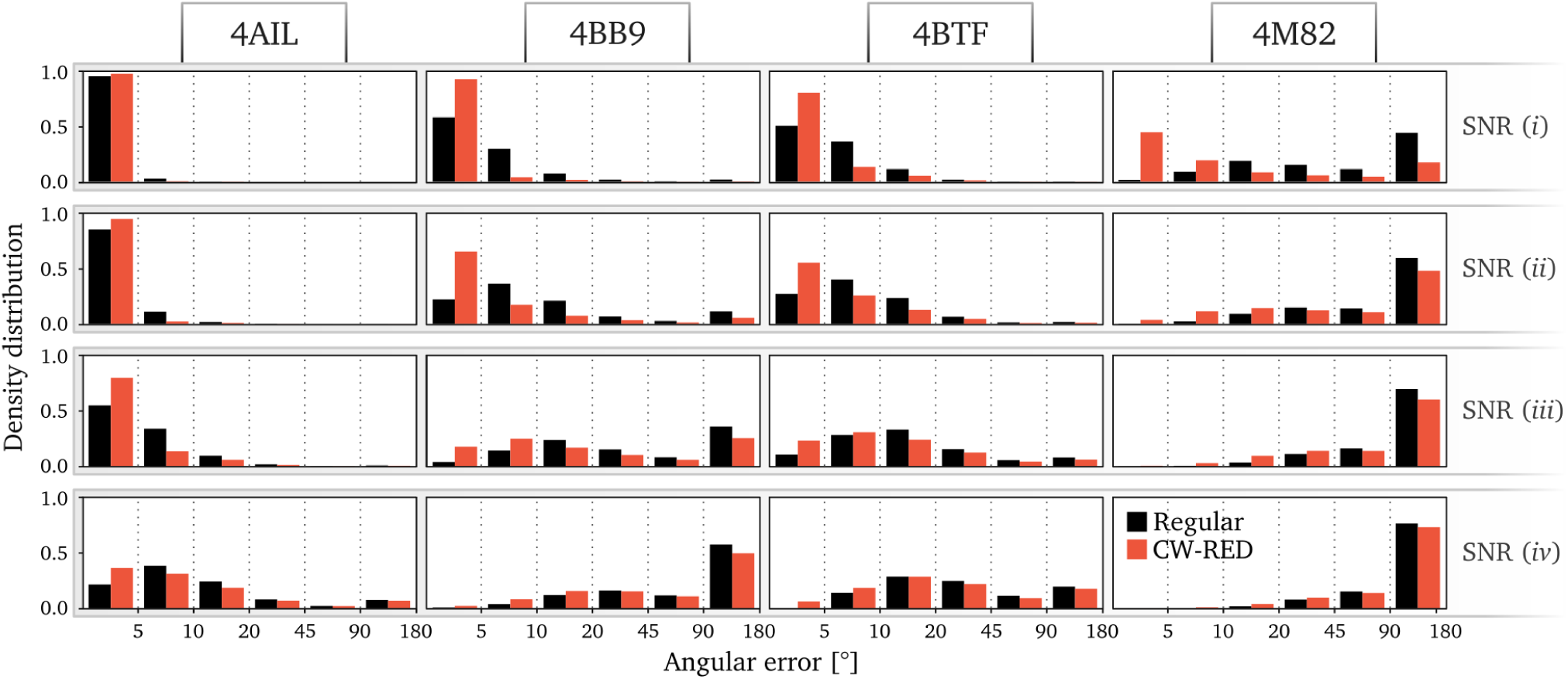
The distribution of angular error of reconstructions at four different SNRs (rows) of four structures (columns). Comparing regular Relion (black) with confidence weighted RED (red). The error is defined as the axis-angle representation difference between the known rotation and the refined angle.

**Figure 6:**
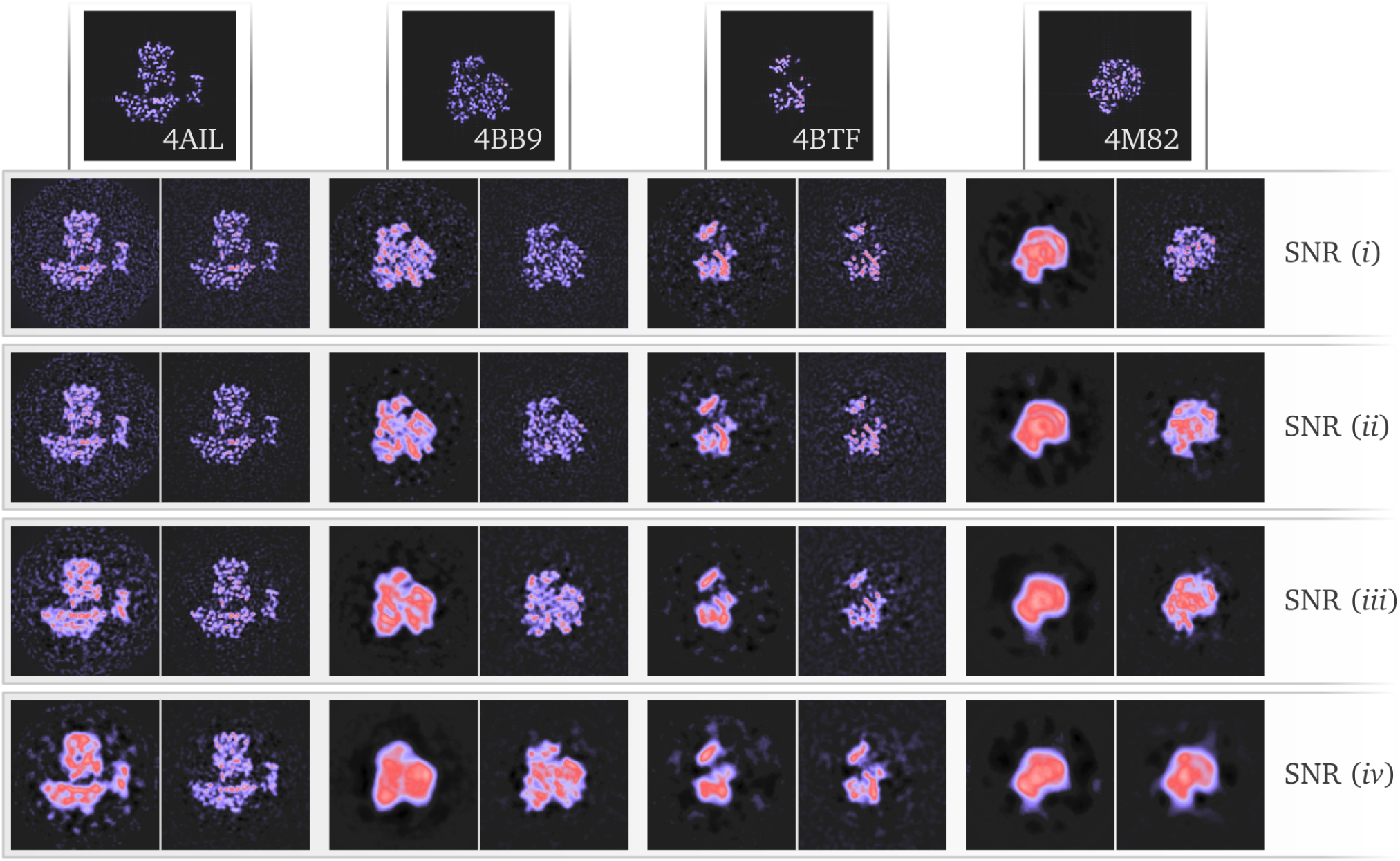
Central slices of the reconstructed maps in the test data set for the four different SNRs. Each pair compares regular Relion (left) with confidence weighted RED (right). Top row shows the ground truth for the map of each column.

**Figure 7:**
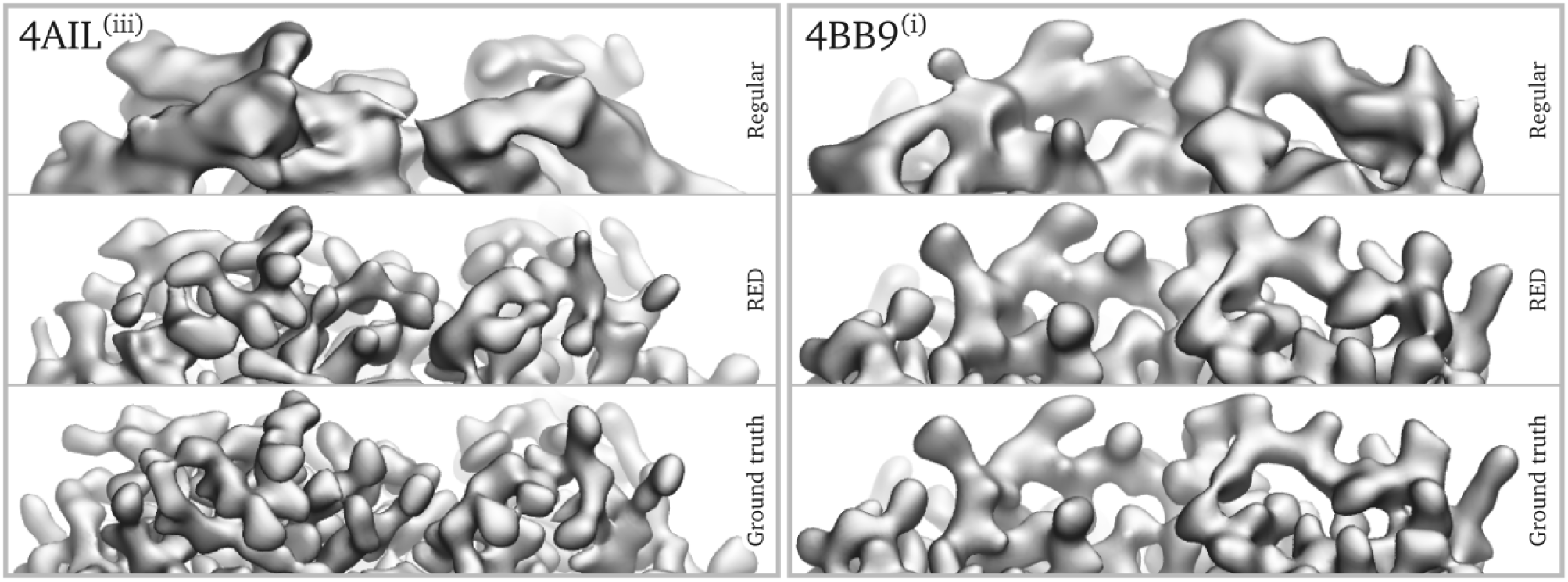
Iso-surface visualisations of the reconstruction results with regular Relion, confidence weighted RED together with ground truth for 4AIL at SNR (*iii*) (left) and 4BB9 at SNR (*i*) (right).

## 5 Discussion

Convolutional neural networks for denoising are typically trained on pairs of noiseless and noisy representations of the signal. It is thus crucial to have access to accurate noiseless ground truth signal, which makes it challenging to apply these networks in areas where such ground truth is impossible or expensive to acquire, such as medical imaging [35]. To circumvent this problem, we used synthetic ground truth cryo-EM reconstructions and used a simplistic physical forward model to generate simulated projection images. In doing so, we explicitly did not aim to obtain a network to denoise reconstructions from experimental cryo-EM data. Rather, we used test data sets that were generated using the same forward model to provide a proof-of-principle that learned priors can improve the reconstruction process. Our results convincingly make this case: the RED approach outperformed standard auto-refinement in Relion in all tests performed.

The standard approach in Relion uses an L2-Tikhonov regularisation on the Fourier components of the reconstruction [2]. In practice, this purely Fourier-based regularisation term is often complemented with an *ad hoc* regularisation in real-space by setting all densities to a constant value in the volume outside a user-specified three-dimensional mask around the reconstruction. Often, such masks are generated after an initial refinement has yielded a preliminary reconstruction. In the tests performed here, no solvent-mask was provided. Thereby, the improvements observed by the RED approach reflect the difference between a purely Fourier-based L2-Thikonov regularisation and learned prior in real-space. The observed differences with masked refinements would be smaller.

A major identified pitfall of this method is the risk of injecting similar features that resemble those of biological macromolecules into both half-maps, which then lead an over-estimation of resolution based on the half-map FSC. Without access to ground truth, the half-map FSC is the single most important measure of quality for the reconstruction, which make this issue a major concern for the usefulness of the method. However, we were able to resolve this issue with an empirical approach that we call confidence-weighted RED. By falling back onto the less informative Gaussian prior at nominal resolutions where the denoiser is known to produce little improvement in the reconstruction, we managed to sufficiently alleviate the problem, while retaining equal overall performance. This approach was here based on performance statistics gathered by applying the denoiser to the training data set, and required no significant additional effort compared to regular RED. To avoid the risk of overestimating the confidence in the denoiser, ideally, another data set should be used instead of the training data set. Furthermore, it is noteworthy that the denoiser performance likely varies over different resolution shells. Therefore, a more detailed investigation into the confidence weighting scheme could lead to further improvements.

We envision multiple other avenues for improving the approach set out in this paper. Firstly, different network architectures may be devised and networks may be trained through alternative loss functions to improve their performance, foremost in the low resolution range. A large variety of successful network architectures have been developed in other areas of image processing, e.g. FFDNet [36] or RCAN [37]. Furthermore, to handle larger reconstructions the denoiser may be trained on patches, rather than on the entire map as done here, and the patches may be combined using a sliding window. This approach is appealing due to the inherent flexibility in terms of particle/box size and memory requirement. Moreover, the use of patches may be intrinsically better suited to deal with the inherent variability in map quality that is caused by different extents of order that exist in many biological macromolecules. However, as networks trained on patches might no longer see the entire box, where a particle at the center is surrounded by solvent, this approach may be less powerful in flattening the solvent. One solution could be the combination of multiple denoisers that are trained in different resolution domains. Confidence weighting may then be employed to each of these denoisers according to their performance in each domain. For instance, a patch-based denoiser dedicated to the high-resolution domain could be combined with a denosier with a global view of the map at low resolution.

Secondly, the networks should be optimised for the denoising of reconstructions from experimental cryo-EM data. Experimental cryo-EM data has different characteristics in both the signal and the noise than the synthetic data used here. Experimental noise is not independent in the pixels, but falls off with higher spatial frequencies. Both the signal and part of the noise are affected by the CTF of the microscope, and in particular the lowest spatial frequencies in experimental cryo-EM reconstructions are poorly modelled by the simplistic forward model of protein atoms in vaccuum used in this paper. Several options exist to generate more realistic pairs of maps for training the denoiser. Refinements with experimental cryo-EM data downloaded from the EMPIAR data base [38] may be used to generate training data. In such a scenario, generating image pairs with ground truth will be difficult, but one could generate intermediate reconstructions from relatively small subsets of the data and provide the high-resolution maps from the complete data sets as substitutes for the ground truth. This is particularly feasible when the networks will only be trained on intermediate-resolution reconstructions (also see below). Alternatively, one could train a generative adversarial networks (GANs) to learn a data-driven forward model from pairs of atomic models and their corresponding noise-free cryo-EM reconstructions, to generate more realistic ground truth maps of disordered regions, e.g. membrane patches. For this purpose, conditional GANs (cGAN) are a particularly suiting candidate, since pairs of atomic model (ground truth) and reconstruction density are available [39]. Similarly, cycle-consistent adversarial networks (CycleGAN) [40] or style transfer GANs [41] may be used to relate the two data domains in cases where pairs are lacking, e.g. for low resolution reconstructions without a matching ground truth or for generalizing the data set to new experimental conditions with few existing examples [42].

Thirdly, the approach described here may be adapted to use optimisation algorithms other than expectation maximisation. A gradient-driven approach with mini-batches would be based on ∇ℒ (*x* | 𝒴) and (15) to inject prior knowledge into the reconstruction more often, which has the potential of improving convergence speed. For this class of algorithms, Adversarial Regularisers might be a more natural candidate compared to RED, since this method models the prior more directly and thus enables for better control of the properties of the generated gradient [43]. Alternatively, algorithms based on ADMM and plug-and-play denoisers are of potential interest for investigation [44, 45].

Finally, an important consideration when employing prior information in solving any inverse problem is that the prior information used in solving the problem can no longer be used for external validation of the results. This is relevant when injecting prior information about biological macromolecules into the cryo-EM reconstruction process. In current cryo-EM approaches such information is not used at all, and the appearance of protein and nucleic-acid-like features in the cryo-EM reconstructions is often implicitly used to confer confidence in the correctness of the maps. However, one could imagine a more informative prior generating reconstructions in which such features are incorrectly “invented” from noise in the maps. Commonly used procedures to prevent overfitting, most notably splitting the data set into two independent halves, do not necessarily protect against interpretation of such false features. Therefore, new validation tools may need to be devised to guard against the interpretation of false features. One option could be to only use information up to a given intermediate resolution when training the denoiser. In that way, the presence of protein or nucleic acid-like features beyond that resolution could still be used to provide confidence in the result. This approach is particularly appealing because the high resolution signal is typically not contributing much to the alignment of the individual noisy projection images anyway [46].

We expect that, with further improvements along the lines discussed above, the approach presented in this paper will result in a tool for structure determination that will outperform the current state-of-the-art, also for experimental cryo-EM data. This has the exciting potential to improve any reconstruction from existing data sets and to expand the applicability of the technique to samples that are currently beyond its scope.

## 6 Acknowledgements

We thank Jake Grimmett and Toby Darling for help with high-performance computing. C.B.S. acknowledges support from the Leverhulme Trust project on ‘Breaking the non-convexity barrier’, the Engineering and Physical Sciences Research Council (EPSRC: EP/M00483X/1, EP/S026045/1 and Centre EP/N014588/1), the RISE projects CHiPS and NoMADS, the Cantab Capital Institute for the Mathematics of Information and the Alan Turing Institute; O.Ö acknowledges support by the Swedish Foundation for Strategic Research grants (AM13-0049 and ID14-0055); S.H.W.S. acknowledges support by the UK Medical Research Council (MC UP A025 1013) and Wave 1 of the UK Research and Innovation (UKRI) Strategic Priorities Fund under the EPSRC (EP/T001569/1), particularly the “AI for Science” theme within that grant and The Alan Turing Institute.

## Appendices

### A Dependent Gaussian noise

SMD describes the results of RED using a white Gaussian noise model with a fixed variance as the underlying assumption for the image distortion. This appendix describes details of how the RED approach was expanded to handle the type of distortions observed in cryo-EM data. In particular, we derive a dynamically adjusted regularization parameter, as described in (15), and a modified training scheme by managing the significantly varying levels of distortion in the data. We show how this leads to a training protocol that minimizes the model mapping to a coarsened ground truth. The aim of this appendix is not to provide a full mathematical proof of (15), but rather present an approach that highlights the assumptions and approximations required to derive it.

We define the random vector 𝕩 ∈ ℂ^*M*^, 𝕩 ∼ *P*, where *P* here is the distribution of undistorted ground truth data. Here, we assume a Gaussian noise model formalized as 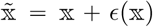, where *ϵ*(𝕩) ∼ 𝒩 (0, *ν*(𝕩)) and *ν* (·): ℂ^*M*^ ⟶ℂ^*M* × *M*^ is a covariance matrix that depends on X to describe the shape of the Gaussian. For the application in this paper this dependency does not have to be explicitly defined, rather an empirical approach is sufficient where the training data set is extended to also include observations of *ν*. In section 3.1 we specify how this was done in practice. One can device an empirical prior distribution that approximates *P* through Kernel Density Estimation (KDE) [47], given a large representative corpus of training data, 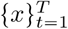, drawn from *P*, and the corresponding 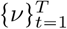, as

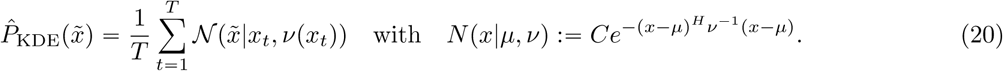

For the continuum limit of KDE, we instead get

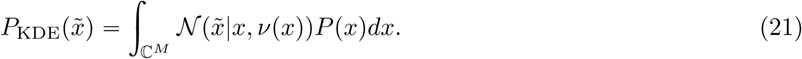

The gradient of the log-KDE-prior then becomes (see Appendix B for the precise definition of ∇)

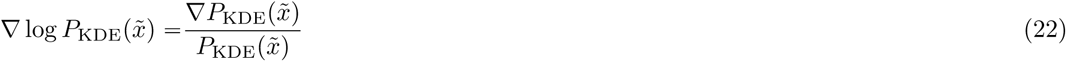

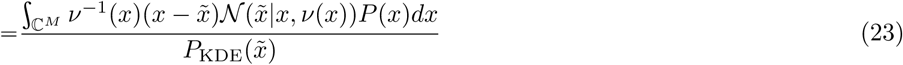

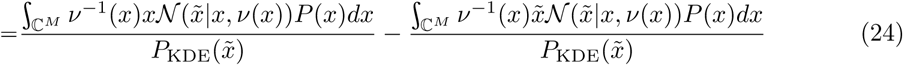

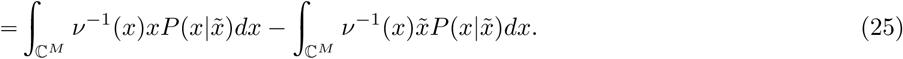

We define an MMSE function as

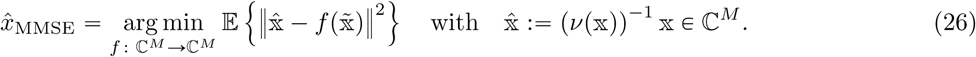

In practice, due to the higher relative error of high-frequency components, 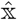 corresponds to a coarsened (low-pass filtered) version of 𝕩. We also define the MMSE of (*ν*(𝕩))^−1^ as

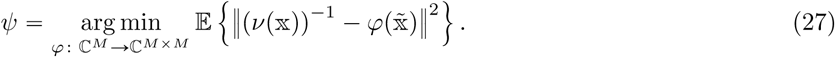

It then holds that

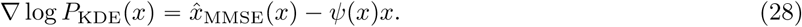

Now, it follows that

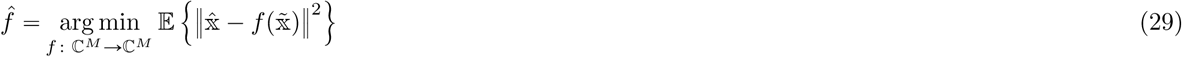

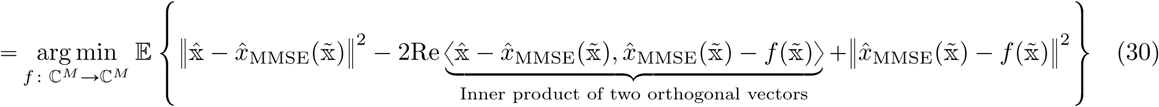

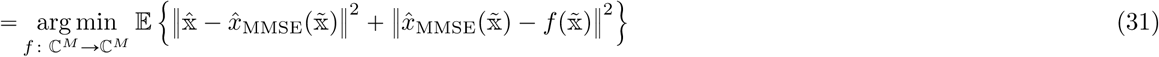

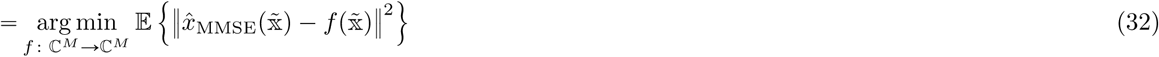

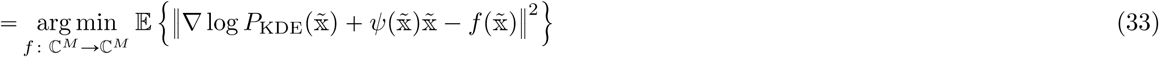

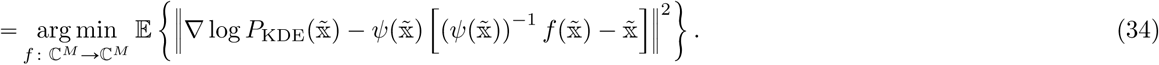

In the above, (31) uses the orthogonality principle and (33) follows from (28). In the particular case of, *f*, in the above, belonging to a class of universal approximator functions, e.g. CNN denoisers, the minimisation (i.e. training) is carried out over the network parameters, *θ*, with pairs of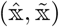. It thus follows that optimizing the weights, *θ*, of such a function, *f*_*θ*_ : ℂ^*M*^ ⟶ ℂ^*M*^, through

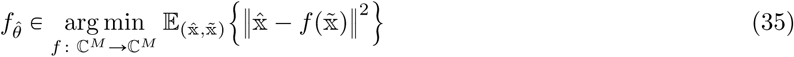

equates matching the gradient of the log-KDE-prior through:

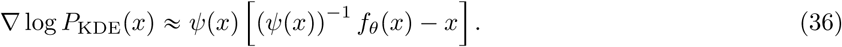

In this paper we assume that *P*_KDE_ is a good enough approximation of *P*, given the training data size and data dimensionality. Conclusively, *f*_*θ*_ is trained to map a noisy data point, 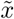, to a weighted combination of low-passed filtered ground truths. We are, however, still required to estimate *ψ*. Ignoring the off-diagonal components of *ψ* and estimating only the diagonal components through (12), i.e. *ψ ≈ τ* ^− 2^ implies that (36) is equivalent to (15).

### B Wirtinger derivatives

To derive eq. (7) one has to differentiate the expression for the posterior probability in (3). In this paper it is expressed in the Fourier domain as a real function with complex input. Here, we define the gradient operator acting on such a function using the formalism of Wirtinger derivatives (see e.g. Appendix A of [48]). More specifically, the Wirtinger derivatives of *f* w.r.t. the complex variables *z*_*i*_*=x*_*i*_*+iy*_*i*_ and 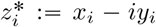 are defined via:

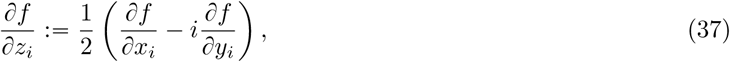

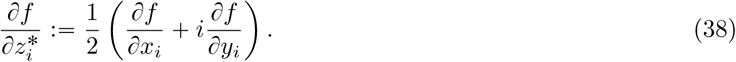

The corresponding gradients can then be defined as

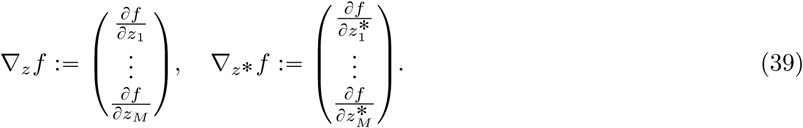

We also use the shorthand notation ∇*f* (*z*) := ∇_*z*\*_ *f* (*z*). Note that *z*_0_ ∈ ℂ^*M*^ is stationary if and only if ∇_*z*_(*z*_0_) = 0 or ∇_*z*\*_ (*z*_0_) = 0. It follows from Appendix A in [48] that for *b, z* ∈ ℂ^*M*^ and *A* ∈ ℂ^*M* × *M*^ :

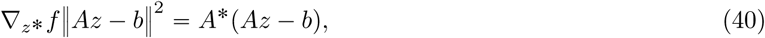

where *A*\* is the adjoint of *A* with respect to the usual innerproduct on ℂ^*M*^. Hence if we take 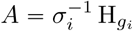 and 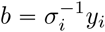 we find

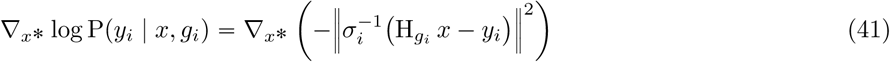

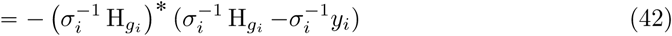

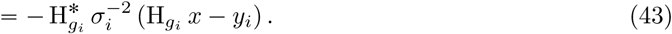

Similar calculations give the gradient of the Gaussian log-prior:

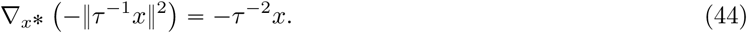

In this paper, to simplify notation outside this appendix, the subscript on∇ is dropped when the differentiation is w.r.t. *x* or any variations thereof.

## Notes

https://github.com/3dem/externprior

### Competing Interest Statement

The authors have declared no competing interest.

### Summary of Updates

Most changes occurred in section 2 (Theory) and the Appendices.

